# Access schedules mediate the impact of high fat diet on ethanol intake and insulin and glucose function in mice

**DOI:** 10.1101/453761

**Authors:** Caitlin R. Coker, Elizabeth A. Aguilar, Angela E. Snyder, Sarah S. Bingaman, Nicholas M. Graziane, Kirsteen N. Browning, Amy C. Arnold, Yuval Silberman

## Abstract

Alcoholism and high fat diet (HFD)-induced obesity individually promote insulin resistance and glucose intolerance in clinical populations, increasing risk for metabolic diseases. Conversely, animal studies, typically utilizing forced/continuous alcohol (EtOH) access, tend to show that EtOH intake mitigates HFD-induced effects on insulin and glucose function, while HFD decreases voluntary EtOH intake in continuous access models. However, the impact of HFD on intermittent EtOH intake and resultant changes to metabolic function are not well characterized. The present studies sought to determine if HFD alters EtOH intake in male C57Bl/6J mice given differing two-bottle choice EtOH access schedules, and to assess resultant impact on insulin sensitivity and glucose tolerance. In the first experiment, mice had Unlimited Access EtOH (UAE)+HFD (n=15; HFD=60% calories from fat, 10% EtOH v/v, *ad libitum*) or UAE+Chow (n=15; control diet=16% calories from fat, *ad libitum*) for 6 weeks. UAE+HFD mice had lower EtOH preference, consumed significantly less EtOH, and were insulin resistant and hyperglycemic compared with UAE+Chow mice. In the second experiment, mice had Limited Access EtOH (LAE, 4 hrs/d; 3 d/wk)+HFD (n=15) or LAE+Chow (n=15) with increasing EtOH concentrations (10%, 15%, 20%). LAE+HFD mice had no difference in total EtOH consumption compared to LAE+Chow mice, but exhibited hyperglycemia, insulin resistance, and glucose intolerance. In the third experiment, mice had intermittent HFD access (single 24 hr session/week) with limited access to EtOH (iHFD-E, 4hrs/d; 4 d/wk) (n=10). iHFD-E mice displayed binge eating behaviors and consumed significantly more EtOH than mice given *ad libitum* chow or HFD, suggesting transfer of binge eating to binge drinking behaviors. Although iHFD-E mice did not have significantly altered body composition, they developed insulin insensitivity and glucose intolerance. These results suggest that access schedules determine the impact of HFD on EtOH consumption and resultant metabolic dysfunction.

## Introduction

Obesity and alcohol use disorder (AUD) are two of the most common chronic conditions in the United States. Clinically, high fat diet (HFD)-induced obesity is associated with insulin resistance, glucose intolerance, and increased risk for developing Type II diabetes (Haslam and James, 2005). Chronic ethanol (EtOH) drinking is also a risk factor for insulin dysfunction (Fan *et al*, 2006; Papachristou *et al*, 2006). Such clinical and epidemiologic findings suggest an overlap in the mechanisms by which HFD and EtOH exposure modulate insulin action and glucose homeostasis and that, together, co-morbid chronic HFD and EtOH intake may increase risk for subsequent disease states such as Type II diabetes (Steiner *et al*, 2015). The majority of epidemiologic evidence, however, suggests that moderate EtOH intake has protective effects on insulin sensitivity (Traversy and Chaput, 2015), but this may only occur in non-obese patients (Yokoyama, 2011). Most preclinical studies also suggest that EtOH consumption mitigates HFD-induced metabolic dysfunction, which may be related to the moderate EtOH intake found in these models (Feng *et al*, 2012; Gelineau *et al*, 2017; Hong *et al*, 2009; Paulson *et al*, 2010). More recent epidemiologic studies, however, have brought these assumptions back up for debate (Griswold *et al*, 2018).

Numerous clinical studies show increased desire, cravings, and intake of high fat foods during and after EtOH drinking episodes (Breslow *et al*, 2013; Caton *et al*, 2004; Piazza-Gardner and Barry, 2014). A similar positive relationship for EtOH-induced increases in HFD intake has been shown in some animal models (Barson *et al*, 2009). While the inverse relationship of HFD exposure stimulating EtOH intake has also been suggested in some animal models (Carrillo *et al*, 2004), the majority of findings indicate that HFD exposure decreases EtOH consumption (Feng *et al*, 2012; Gelineau *et al*, 2017; Sirohi *et al*, 2017a, 2017b). The majority of these previous studies, however, have examined EtOH intake in the face of HFD access without specific focus on metabolic function, or examined metabolic effects of EtOH and HFD without taking intake behaviors into account. While the studies above and others (Guo *et al*, 2018) have greatly advanced our understanding of the impact that HFD and EtOH have on metabolic and end-organ function, these models typically do not replicate the escalation of drinking behaviors common to human AUD.

It is now well characterized that limited access scheduling increases EtOH intake in animal models in an escalating fashion akin to human AUD development (Melendez, 2011). The impact of HFD on this type of scheduled EtOH access, however, has not been examined. Intermittent HFD access has been shown to induce binge eating behaviors in mice (Czyzyk *et al*, 2010; Hardaway *et al*, 2016). Since acute HFD can increase EtOH intake (Carrillo *et al*, 2004), and vice versa (Barson *et al*, 2009), it stands to reason that repeated acute HFD access (i.e. intermittent HFD) might lead to increased escalation of EtOH intake under intermittent EtOH access conditions. Therefore, the overall goals of these studies were to determine: (1) if HFD alters EtOH intake in mice consuming EtOH with limited or unlimited access schedules; and (2) how such access schedules modulate the interaction of HFD and EtOH on metabolic function in male C57Bl/6J mice. Our findings suggest that continuous HFD reduces EtOH intake when EtOH is freely available, but that HFD does not alter EtOH intake when access to EtOH is limited. Furthermore, intermittent access to HFD significantly increases EtOH intake when EtOH is also available on an intermittent schedule. Contrary to previous work, our study suggests that moderate intake of freely-available EtOH does not mitigate the ability of HFD to promote insulin resistance. In HFD-fed mice, higher levels of EtOH intake in the limited access models failed to improve insulin sensitivity and worsened glucose tolerance, suggesting scheduled HFD and EtOH intake may interact to disrupt insulin action and glucose homeostasis.

## Methods

### Animals

Six-week old male C57Bl/6J mice were purchased from The Jackson Laboratory (stock # 000664). Upon arrival, mice were individually housed and given standard chow diet for a four-day acclimation period. Mice were weight matched and separated into groups as described below. All mice were kept in a temperature- and humidity-controlled room on a 12-hour light/dark cycle. All procedures were approved by the Institutional Animal Care and Use Committee at Penn State University College of Medicine (Hershey, PA).

### EtOH Ramp

All mice underwent an EtOH-ramp initiation period to avoid potential confounds of EtOH taste aversion. The EtOH two-bottle choice ramp procedure consisted of home-cage 24-hour access to a bottle containing tap water and another bottle containing 3% EtOH for 48 hours, 7% EtOH for 72 hours, and 10% EtOH for 72 hours (all EtOH concentrations are vol/vol in tap water). Water and EtOH solutions were administered via inverted 50 mL conical tubes (Fisher) and sealed with a rubber stopper (#5.5, Fisher) containing a 2-inch stainless-steel straight sipper tube (Ancare). EtOH solution was made using ethyl alcohol (190 proof, PHARMCO-AAPER) diluted in tap water.

### Experiment 1: Effects of ad libitum diet on unlimited access to EtOH (UAE model)

To examine effects of continuous two-bottle choice EtOH drinking in the presence of *ad libitum* HFD (60% calories from fat, Bioserv F3282) or control diet (16% calories from fat, Bioserv F4031), mice were weight matched and randomly assigned into an unlimited 24-hour access to EtOH group receiving either HFD (UAE+HFD; n=15) or control diet (UAE+Chow; n=15). Following a 2-week HFD exposure period and the EtOH-ramp initiation period, UAE+HFD and UAE+Chow groups had home-cage 24-hour access to two-bottle choice of tap water and 10% EtOH (vol/vol in water) throughout the remainder of the experiment. Body mass, EtOH and water intake, and EtOH preference were assessed every 24 hours for six weeks. Summary of timeline provided in Fig 1.

**Fig 1.**
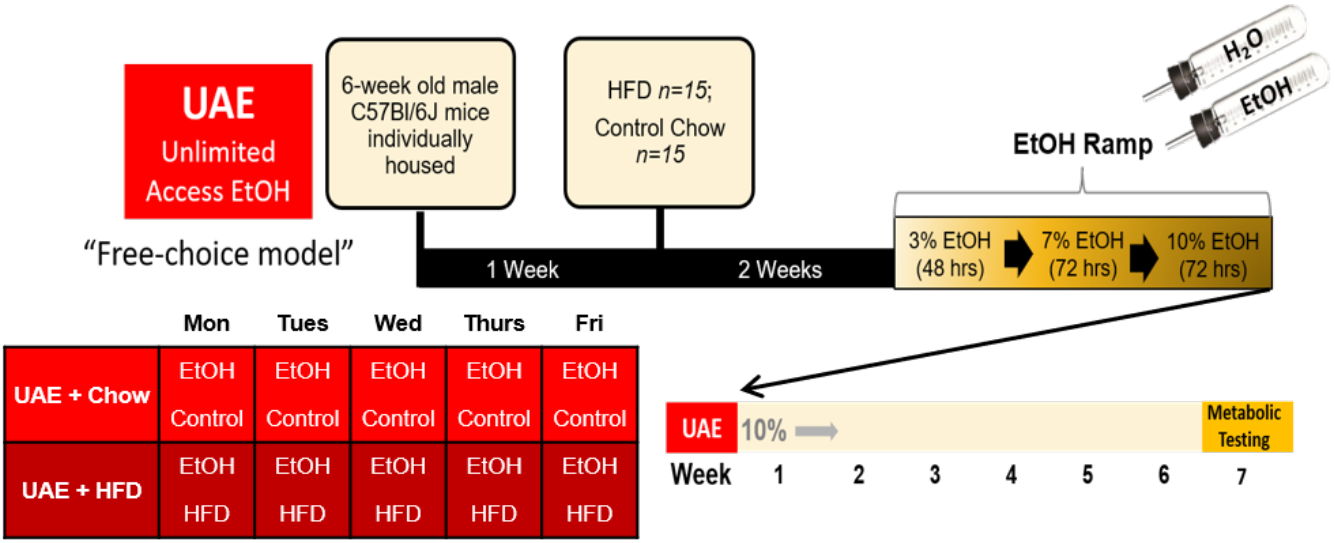
Summary of free-choice, Unlimited Access Ethanol (UAE) timeline.

### Experiment 2: Effects of ad libitum diet on limited access to EtOH (LAE model)

To examine effects of limited two-bottle choice EtOH drinking in the presence of *ad libitum* HFD or control diet, mice were weight matched and randomly assigned to groups receiving either HFD (LAE+HFD; n=15) or control diet (LAE+Chow; n=15). Diets are the same as described above, except that mice had an initial 48-hour HFD access period (or control Chow) prior to the EtOH-ramp. Following the EtOH-ramp initiation period, appropriate diets were then provided *ad libitum* for the remainder of the study. LAE+HFD and LAE+Chow groups had home-cage two-bottle choice of tap water and EtOH (vol/vol in water) limited to 4-hour access periods on Monday, Wednesday, and Friday beginning at 10am and ending at 2pm. These groups had access to 10% EtOH for three weeks, followed by 15% EtOH for two weeks, and 20% EtOH for two weeks. Body mass, EtOH and water intake, and EtOH preference were assessed after each drinking session.

### Experiment 3: Effects of intermittent HFD access on limited access to EtOH (iHFD-E model)

To examine effects of limited two-bottle choice EtOH drinking in the presence of intermittent HFD, mice were weight matched and randomly assigned to groups receiving either intermittent HFD (iHFD-E; n=10), *ad libitum* HFD (HFD-E; n=10) or *ad libitum* control chow diet (Chow-E; n=10). iHFD-E mice had a single 24-hour HFD access period per week (Czyzyk *et al*, 2010). After an initial 48-hour HFD access period (or control Chow) and EtOH-ramp initiation period, all groups began their appropriate diet regimens and began home-cage two-bottle choice of tap water and EtOH (vol/vol in water) limited to 4-hour access periods on Tuesday, Wednesday, Thursday, and Friday beginning at 10am and ending at 2pm. These groups had access to 10% EtOH for three weeks, followed by 15% EtOH for two weeks, and 20% EtOH for two weeks. Body mass, EtOH and water intake, and EtOH preference were assessed after each drinking session.

### Metabolic Testing

Once EtOH intake studies concluded, mice underwent standardized insulin tolerance (ITT) and glucose tolerance (GTT) tests. For the ITT, mice were fasted for four hours and then injected intraperitoneally with insulin (0.75 units/kg of regular U-100 insulin in 1x PBS; Novolin). A tail vein blood sample was taken at baseline (immediately prior to insulin injection) and at 15, 30, 60, 90, and 120 minutes post-insulin injection to measure blood glucose with a glucometer (Prodigy AutoCode). For the GTT, mice were fasted overnight and then injected intraperitoneally with dextrose (2 g/kg of 50% dextrose). Blood glucose was measured at baseline (immediately prior to dextrose injection) and at 15, 30, 60, 90, and 120 minutes post-dextrose injection. Body composition (fat, lean, and fluid masses) was measured in conscious mice using a quantitative nuclear magnetic analyzer (Bruker Minispec). Age-and weight-matched mice receiving HFD (n=10) or control diet (n=10) for similar periods but without EtOH exposure were used as controls.

### Statistics

Two-way repeated measures ANOVA was used to compare the main effects of diets and number of drinking sessions on EtOH intake, and their interaction over the study time course. One-way ANOVA was used to compare end of study metabolic metrics between diet+EtOH groups versus diet exposed control groups not receiving EtOH. When applicable, one-way ANOVA or unpaired t-tests were used to compare end of study EtOH consumption metrics between diet+EtOH groups. All data are represented as mean±standard error of the mean (SEM).

## Results

### Experiment 1

We determined effects of *ad libitum* HFD or chow diet access on body mass and EtOH intake parameters in the UAE model, which had continuous free-access to 10% EtOH and water. Two-way ANOVA indicated that UAE+HFD mice had significantly higher body mass than UAE+Chow mice over the course of the study (Diet: F_(1,18)_=44.91, p<0.001; EtOH exposure sessions: F_(29,522)_=163.1, p<0.001; Interaction: F_(29,522)_=76.23, p<0.001; Fig. 4A). UAE+HFD mice consumed significantly less EtOH than UAE+Chow mice (Diet: F_(1,18)_=22.22, p<0.001; EtOH exposure sessions: F_(24,432)_=3.927, p<0.001; Interaction: F_(24,432)_=1.250, p=0.194; Fig. 4B). Given that the large differences in body mass could skew evaluation of g/kg measurements, we also assessed total grams of EtOH (g/EtOH) consumed per group per day. Two-way ANOVA confirmed a reduction in g/EtOH by HFD (Diet: F_(1,18)_= 8.80, p<0.01; EtOH exposure sessions: F_(24,432_)=4.01, p<0.0001; Interaction: F_(24,432)_= 0.9, p=0.597; Fig. 4C). Cumulative g/EtOH consumption was higher in UAE+Chow mice compared to UAE+HFD mice (3.11±0.36 vs 1.89±0.20 g/EtOH, respectively, t=2.966, df=18, p=0.008). UAE+HFD mice also had lower preference towards EtOH (Diet: F_(1,18)_=4.700, p=0.044; EtOH exposure sessions: F_(25,450)_=4.872, p<0.001; Interaction: F_(25,450)_=0.8202; p=0.717; Fig. 4D). These findings indicate that *ad libitum* HFD access reduces EtOH intake in unlimited access “free-choice” model.

At the end of the study, mice underwent metabolic testing. Metabolic data were compared to weight- and age-matched mice given HFD or chow diet without EtOH access. One-way ANOVA (F_(3,38)_=26.02, p<0.001; Fig. 5A) indicated that body mass was similarly elevated in EtOH-naïve HFD (37.9±1.1 g) and UAE+HFD exposed (42.1±2.1 g) mice when compared to EtOH-naïve Chow (28.5±0.5 g) or UAE+Chow (28.4±0.3 g) mice. One-way ANOVA indicated that adiposity was significantly altered by EtOH consumption and that this was further impacted by HFD consumption (F_(3,38)_=27.02, p<0.0001; Fig 5B). Bonferroni’s post-hoc analysis showed a significant increase in adiposity in the UAE+Chow and EtOH-naïve HFD mice compared to EtOH-naïve Chow mice and a further increase in adiposity in UAE+HFD mice compared to all other groups (Fig 5B). One-way ANOVA with Bonferroni’s post hoc analysis showed that lean mass was reduced in UAE+Chow, UAE+HFD, and EtOH-naïve HFD mice compared to EtOH-naïve Chow mice, but this reduction in lean mass was not as pronounced in UAE+Chow mice(F_(3,38)_=25.40, p<0.001; Fig. 5C). One-way ANOVA further showed that HFD can increase fluid mass but that this did not appear to be altered by EtOH consumption (F_(3,38)_=7.92, p<0.001; Fig. 5D). Overall, these data indicate that EtOH consumption may have subtle effects on HFD-induced changes in body composition in this model, with most pronounced effects on adiposity.

HFD-induced increases in body mass and adiposity are accompanied by insulin resistance in mouse models, and previous research indicates that EtOH consumption may mitigate these effects. We therefore performed ITT in the UAE mice and compared these results to the same age- and weight-matched controls as utilized in Fig 5. One-way ANOVA (F_(3,38)_=3.305, p=0.030; Fig. 6A) followed by Bonferroni’s post-hoc analysis indicated a significant increase in 4-hour fasting glucose levels in HFD (209±6 mg/dl) compared to Chow (166±7 mg/dl) mice, with no statistically significant differences between UAE+HFD (202±14 mg/dl) or UAE+Chow groups (182±12 mg/dl) compared to controls. Fig. 6B shows the time course of change in blood glucose levels following intraperitoneal insulin injection; data are normalized to baseline to account for differences in fasting glucose levels among groups. Fig. 6C shows the area under the curve for changes in blood glucose levels over time in response to insulin administration, with a more negative value indicating better insulin sensitivity. The ITT area under the curve indicated reduced insulin sensitivity in EtOH-naïve HFD mice (975±968 glucose mg/dl*min) compared to Chow mice (−4321±852 glucose mg/dl*min), with no effect of EtOH consumption on HFD-induced insulin resistance in UAE+HFD mice (−1282±1091 glucose mg/dl*min) (F_(3,38)_=4.624, p=0.008; Fig. 6C). UAE+Chow mice were not significantly different from EtOH-naïve Chow mice. Contrary to many previous findings, these results indicate that free-access EtOH consumption does not improve insulin sensitivity in HFD exposed mice.

### Experiment 2

The above findings indicate that HFD decreases EtOH intake when provided in an unlimited “continuous-access” procedure. Previous research indicates that limiting EtOH access in an every-other-day intermittent access model increases EtOH intake in rodents compared to continuous access (Melendez, 2011). Whether HFD alters intermittent EtOH intake remains unclear. Therefore, we placed weight-matched mice in a limited access EtOH procedure (LAE; 4hr/day, Monday, Wednesday, Friday every week, see Fig. 2) and gave mice *ad libitum* HFD or chow. Two-way ANOVA indicated LAE+HFD mice gained significantly more body mass than LAE+Chow mice over the course of the study (Diet: F_(1,28)_=60.72, p<0.001; EtOH exposure sessions: F_(20,560)_=522.7, p<0.001; Interaction: F_(20,560)_=180.8, p<0.001; Fig. 7A). LAE+HFD mice had significantly lower EtOH g/kg levels than the LAE+Chow group (Diet: F_(1,28)_=18.37, p<0.001; EtOH exposure sessions: F_(20,560)_=52.83, p<0.001; Interaction: F_(20,560)_=5.734, p<0.001; Fig. 7B). Given that the large differences in body mass could skew evaluation of g/kg measurements, we also assessed total grams of EtOH (g/EtOH) consumed. Two-way ANOVA indicated a significant effect of exposure session (EtOH exposure sessions: F_(20,560)_=65.20, p<0.001) with no statistically significant effect of diet on g/EtOH consumed (Diet: F_(1,28)_=3.902, p=0.058) but a significant interaction between the two variables (Interaction: F_(20,560)_=1.645, p=0.039; Fig. 7C). Further, there was no significant effect of diet on cumulative g/EtOH consumed over the course of the study between LAE+Chow vs LAE+HFD mice (1.91±0.13 vs 1.53±0.15 g/EtOH, respectively, t=1.975, df=28, p=0.0582; Fig. 7C **inset**). For EtOH preference, two-way ANOVA indicated significant main effects of diet (F_(1,28)_=18.85, p<0.001) and EtOH exposure sessions (F_(20,560)_=5.967, p<0.001), with no significant interaction (F_(20,560)_=1.376, p=0.127; Fig. 7D).

**Fig 2.**
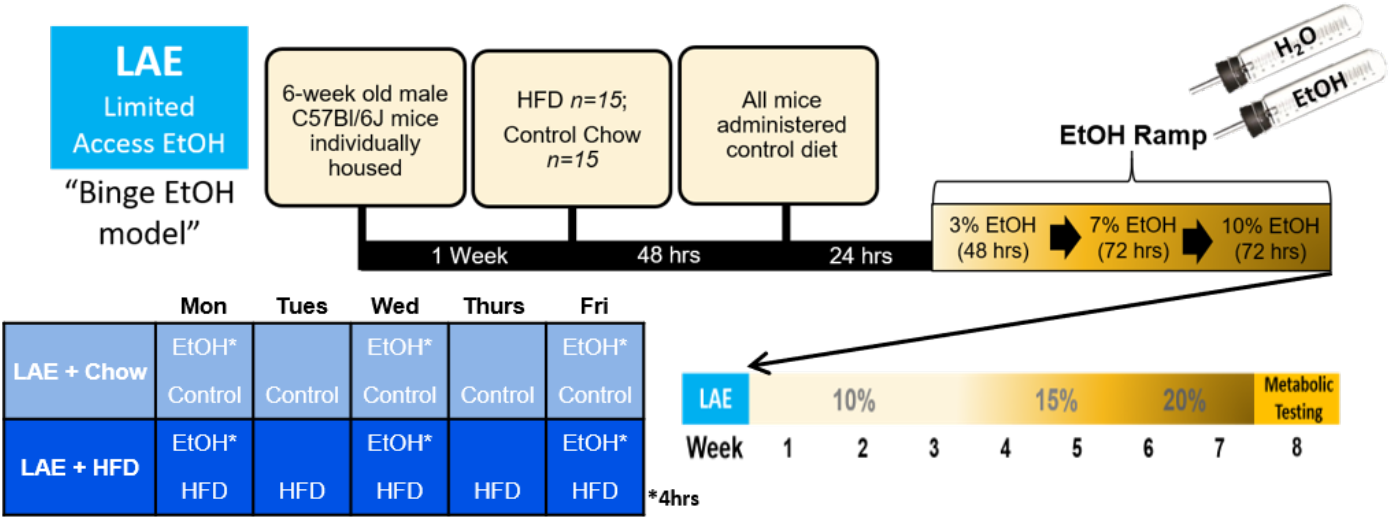
Summary of Limited Access Ethanol (LAE) timeline.

For metabolic testing, results from LAE mice were compared to the same EtOH-naïve HFD and Chow control mice utilized in the previous experiment. One-way ANOVA showed body mass was elevated in both EtOH-naïve HFD (37.9±1.1 g) and LAE+HFD (43.8±1.1 g) mice when compared to LAE+Chow (28.1±0.6 g) or chow-fed EtOH-naïve mice (28.5±0.5 g) (F_(3,46)_=77.62, p<0.001; Fig. 8A). In contrast to the UAE model, limited EtOH access in the LAE+HFD mice led to a significantly increased body mass compared to EtOH-naïve HFD mice. Since EtOH intake did not alter body mass in LAE+Chow mice compared to EtOH-naïve Chow mice, these findings suggest a potential synergistic interaction between HFD and EtOH intake on body mass. One-way ANOVA indicated that HFD and EtOH consumption significantly altered adiposity (F_(3,46)_=288.9, p<0.001; Fig. 8B), lean mass (F_(3,46)_=280.6, p<0.001; Fig. 8C), and fluid mass (F_(3,46)_=87.68, p<0.001; Fig. 8D). For each measure tested, potential synergistic interactions between HFD and EtOH on body mass were further reflected by significantly increased adiposity and fluid mass in LAE+HFD vs EtOH-naïve HFD mice and decreased lean mass between these two groups. LAE+Chow mice were not significantly different from EtOH-naïve Chow mice on any of these measures.

For the ITT, one-way ANOVA (F_(3,46)_=31.70, p<0.001; Fig. 9A) followed by Bonferroni’s post-hoc test indicated 4-hour fasting glucose was elevated in the EtOH-naïve HFD (209±6 mg/dl) and LAE+HFD (250±8 mg/dl) mice compared to EtOH-naïve Chow (166±7 mg/dl) and LAE+Chow (173±7 mg/dl) mice. Similar to body composition data, fasting glucose was significantly higher in the LAE+HFD mice compared to EtOH-naïve HFD mice although LAE+Chow mice were not significantly different from EtOH-naïve Chow mice. One-way ANOVA (F_(3,44)_=5.229, p=0.004; Fig. 9B,C) indicated that insulin sensitivity was reduced in the EtOH-naïve HFD group (975±968 glucose mg/dl*min) compared to EtOH-naïve Chow mice (−4321±852 glucose mg/dl*min). There was no significant difference in insulin sensitivity between LAE+Chow (−3583±460 glucose mg/dl*min) and EtOH-naïve Chow mice. Surprisingly, although there was a significant increase in resting glucose in LAE+HFD mice, there was no significant difference in insulin sensitivity in these mice (−1885±1281 glucose mg/dl*min) compared to any other group.

Given differences in fasting glucose levels between groups, we next performed GTT to determine the impact of HFD and EtOH on the ability to dissipate changes in blood glucose in response to a glucose load. One-way ANOVA indicated that 12-hour fasting glucose levels were elevated in HFD compared to chow-fed mice (F_(3,46)_=22.36, p<0.001; Fig. 9D), but EtOH intake did not alter HFD-induced changes as LAE+HFD (197±7 mg/dl) was not statistically different from EtOH-naïve HFD mice (205±9 mg/dl). The increase in blood glucose levels in response to exogenous dextrose administration over the 120-minute study period is shown in Fig. 9E. These data were summarized as an area under the curve in Fig. 9F, with a more positive value indicating glucose intolerance. One-way ANOVA (F_(3,46)_=30.09, p<0.001; Fig. 9E,F) indicated a significant increase in the GTT area under the curve in LAE+HFD mice (38047±2397 glucose mg/dl*min) and EtOH-naïve HFD mice (36322±2421 glucose mg/dl*min) compared to LAE+Chow (19028±1493 glucose mg/dl*min) and EtOH-naïve Chow (17699±938 glucose mg/dl*min) mice.

### Experiment 3

Since limiting access to HFD can induce binge eating behaviors (Czyzyk *et al*, 2010; Hardaway *et al*, 2016), we hypothesized that such binge intake behaviors toward food would transfer to EtOH intake behaviors to increase EtOH intake. We therefore sought to examine the effects of intermittent HFD access on limited access two-bottle choice EtOH drinking. We placed weight-matched mice in a limited access EtOH procedure (4hr/day, Tuesday, Wednesday, Thursday, Friday every week, see Fig 3) and gave mice intermittent access to HFD (a single 24hr period/week; iHFD-E), *ad libitum* chow (Chow-E), or *ad libitum* HFD (HFD-E). Two-way ANOVA indicated HFD-E mice gained significantly more body mass than Chow-E or iHFD-E mice over the course of the study (Diet: F_(2,27)_=81.04, p<0.001; EtOH exposure sessions: F_(34,918)_=363.4, p<0.001; Interaction: F_(68,918)_=89.98, p<0.001; Fig. 10A). Interestingly, although weight gain was predominantly linear in HFD-E and Chow-E mice, iHFD-E mice showed weight cycling, indicating a binge-like eating pattern where most of their food intake each week occurred on the HFD exposure day with reduced food intake the rest of the week. Two-way ANOVA indicated that iHFD-E mice had significantly higher g/kg levels (Diet: F_(2,27)_=33.08, p<0.001; EtOH exposure sessions: F_(27,729)_=25.02, p<0.001, Interaction: F_(54,729)_=12.5, p<0.001) and higher g/EtOH consumed (Diet: F_(2,27)_=26.55, p<0.001; EtOH exposure sessions: F_(27,729)_=25.12, p<0.001; Interaction: F_(54,729)_=11.17, p<0.001) than Chow-E or HFD-E (Fig. 10B,C). Total EtOH consumption over the course of the study was also significantly higher in iHFD-E mice than Chow-E or HFD-E mice (One-way ANOVA, F_(2,27)_=26.55, p<0.001; Fig. 10C **inset**) while post-hoc analysis showed no significant difference in total EtOH consumption between Chow-E and HFD-E mice. Two-way ANOVA also indicated that although there was no significant effect of diet on EtOH preference (F_(2,27)_=2.663, p=0.088), there was a significant effect of EtOH exposure sessions (F_(27,729)_=2.938, p<0.001) and interaction (F_(54,729)_=3.295, p<0.001; Fig. 10D).

**Fig 3.**
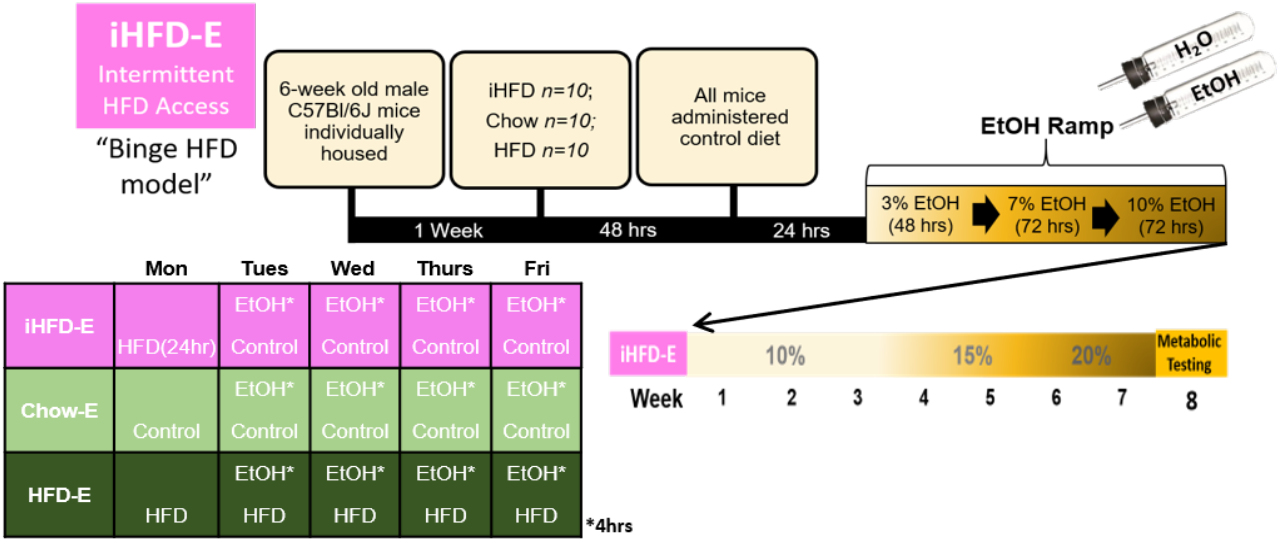
Summary of intermittent access to HFD with limited access to EtOH (iHFD-E) timeline.

For metabolic testing, results from iHFD-E, Chow-E, and HFD-E mice were compared to the same EtOH-naïve HFD and Chow control mice utilized in the previous experiments. One-way ANOVA showed body mass was elevated in both EtOH-naïve HFD (37.9±1.1 g) and HFD-E (42.7±1.0 g) mice when compared to Chow-E (27.0±0.4 g), iHFD-E (29.4±0.8 g), or EtOH-naïve Chow mice (28.5±0.5 g) (F_(4,45)_=70.28, p<0.001; Fig. 11A). Body mass did not differ between Chow-E, iHFD-E, or EtOH-naïve Chow mice, but similar to findings in Experiment 2, HFD-E mice gained significantly more body mass than EtOH-naïve HFD mice. Overall, one-way ANOVA indicated a significant change of adiposity (F_(4,45)_=96.79, p<0.001; Fig. 11B), lean mass (F_(4,45)_=105.6, p<0.001; Fig. 11C), and fluid mass (F_(4,45)_=94.85, p<0.001; Fig. 11D). For individual groups, there appeared to be no significant effect of EtOH consumption in the Chow-E group on any body composition measurement compared to EtOH-naïve Chow mice. EtOH-naïve HFD mice had increased body mass, adiposity, and fluid mass, and reduced lean mass (compared to Chow mice), effects that were exacerbated by EtOH consumption in the HFD-E mice. iHFD-E mice had a reduction in lean mass compared to Chow mice, but otherwise had no significant changes to body mass, adiposity, or fluid mass.

**Fig 4.**
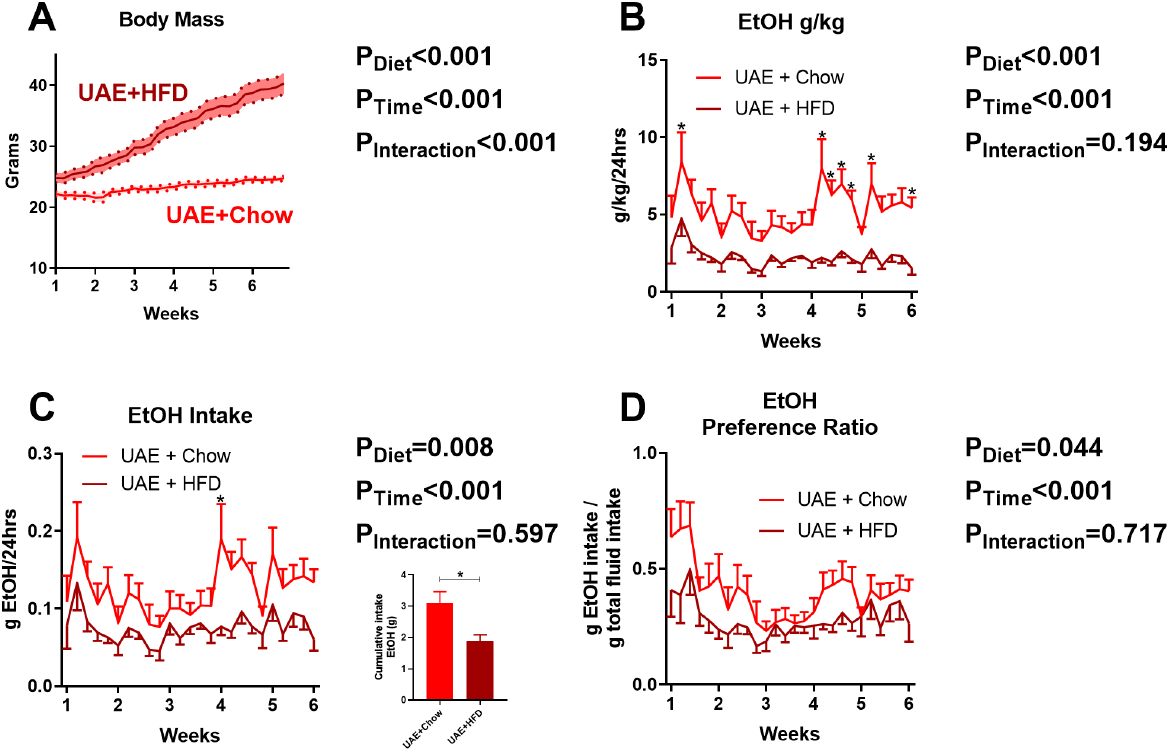
*Ad libitum* HFD access reduces EtOH intake in unlimited access “freechoice” UAE model. **A)** Time course of body mass changes by group during drinking period (n=15/group). Dark line indicates mean, shaded area with dots indicates range of standard error of the mean. **B-D)** HFD significantly reduces EtOH g/kg/24hrs, total EtOH g consumed/24hrs, and EtOH preference. Line indicates mean, error bars indicates standard error of mean. * indicates significant effect of diet as determined by two-way ANOVA; p<0.05.

**Fig 5.**
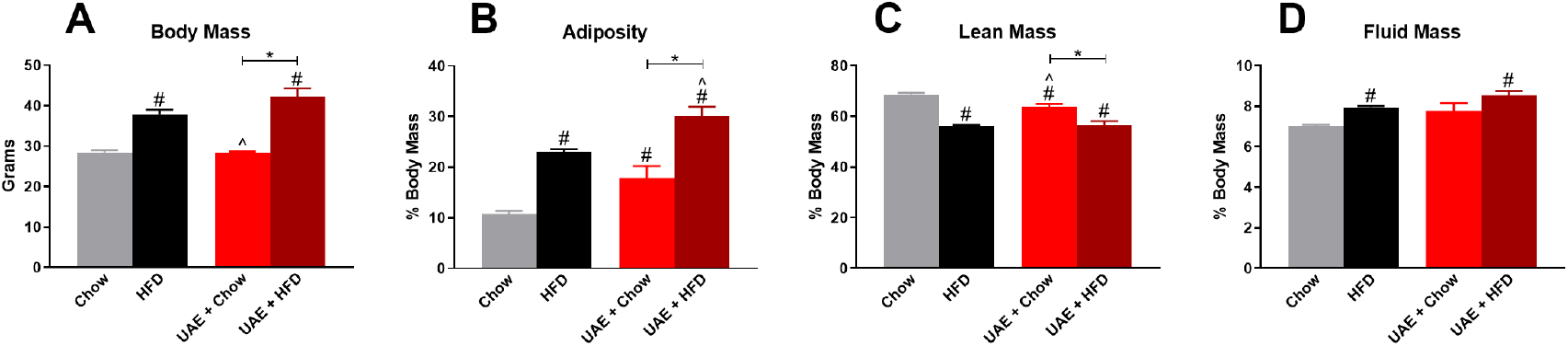
Free access EtOH consumption alters HFD-induced changes in adiposity with no change in body mass, lean mass, or fluid mass. Body composition data are compared to chow- and HFD-fed EtOH-naïve mice. **A)** Body mass of control mice and LAE mice prior to metabolic testing. Percent body mass of **B)** adiposity, **C)** lean mass, and **D)** fluid mass. * indicates significant difference between indicated groups, # indicates significant difference from Chow, ^ indicates significant difference from HFD, as determined by one-way ANOVA with Bonferroni’s post-hoc test,

**Fig 6.**
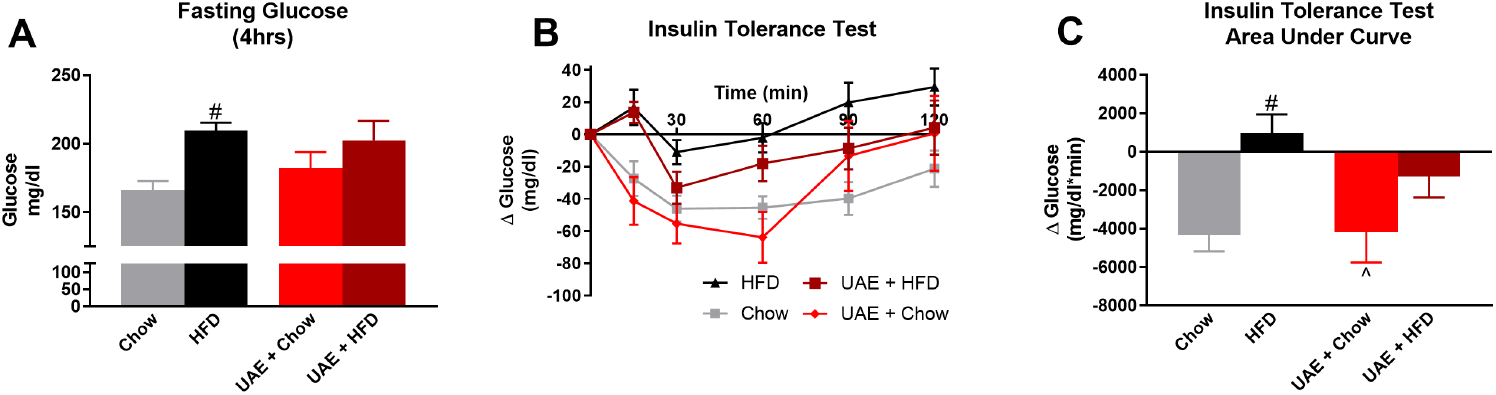
Moderate EtOH consumption does not alter HFD-induced insulin resistance. **A)** 4-hour fasting glucose levels prior to insulin tolerance test. **B)** Change in blood glucose levels over time following insulin injection; data are normalized to 0 at baseline. **C)** Area under the curve for change in blood glucose levels during insulin tolerance test. One-way ANOVA; * indicates significant difference between indicated groups, # indicates significant difference from Chow, ^ indicates significant difference from HFD, as determined by one-way ANOVA with Bonferroni’s post-hoc test, p<0.05.

For the ITT, one-way ANOVA (F_(4,45)_=38.65, p<0.001; Fig. 12A) followed by Bonferroni’s post-hoc test indicated that 4-hour fasting glucose was elevated in the EtOH-naïve HFD (209±6 mg/dl) mice compared to Chow-E (162±5 mg/dl), iHFD-E (157±10 mg/dl), or EtOH-naïve Chow mice (166±7 mg/dl), with an even greater increase in HFD-E mice (271±10 mg/dl). One-way ANOVA (F_(4,45)_=6.9, p<0.001; Fig. 12B,C) indicated that insulin sensitivity was reduced in the EtOH-naïve HFD group (975±968 glucose mg/dl*min) compared to EtOH-naïve Chow mice (−4321±852 glucose mg/dl*min), Chow-E mice (−3823±809 glucose mg/dl*min), and HFD-E mice (−3513±882 glucose mg/dl*min), with a trend for reduced insulin sensitivity in iHFD-E mice (−1699±559 glucose mg/dl*min). For the GTT, one-way ANOVA (F_(4,45)_=6.637, p<0.001; Fig. 12D) indicated that 12-hour fasting glucose levels were elevated in EtOH-naïve HFD mice (205±9 mg/dl) and HFD-E mice (186±11 mg/dl) compared to EtOH-naïve Chow mice (138±7 mg/dl), but iHFD-E mice (179±16 mg/dl) were not statistically different from any other group. One-way ANOVA (F_(4,45)_=17.07, p<0.001; Fig. 12E,F) indicated a significant increase in the GTT area under the curve in HFD-E (37634±2428 glucose mg/dl*min) compared to Chow-E mice (22871±1827 glucose mg/dl*min), iHFD-E mice (28952±2342 glucose mg/dl*min), and EtOH-naïve Chow mice (17699±938 glucose mg/dl*min). There was also a significant increase in GTT AUC in iHFD-E mice (28952±2342 glucose mg/dl*min) compared to EtOH-naïve Chow mice (17699±938 glucose mg/dl*min).

**Fig 7.**
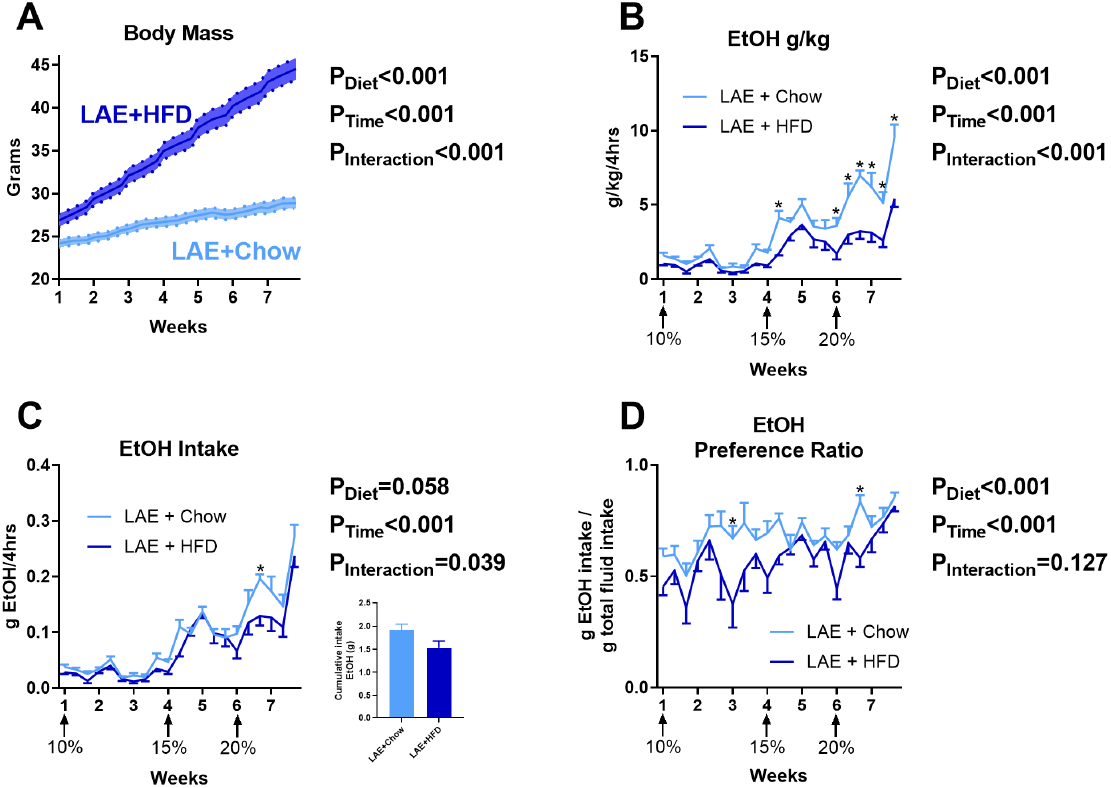
HFD effects on EtOH intake in the limited access “LAE” model. **A)** Time course of body mass changes by group during drinking period (n=15/group). HFD decreased g/kg intake measure **(B)** but does not significantly alter total g/EtOH consumed over the course of the study **(C)** and modestly alters preference **(D)**. * indicates significant differences between groups on individual EtOH access sessions as determined by Bonferroni post-hoc analysis; p<0.05.

**Fig 8.**
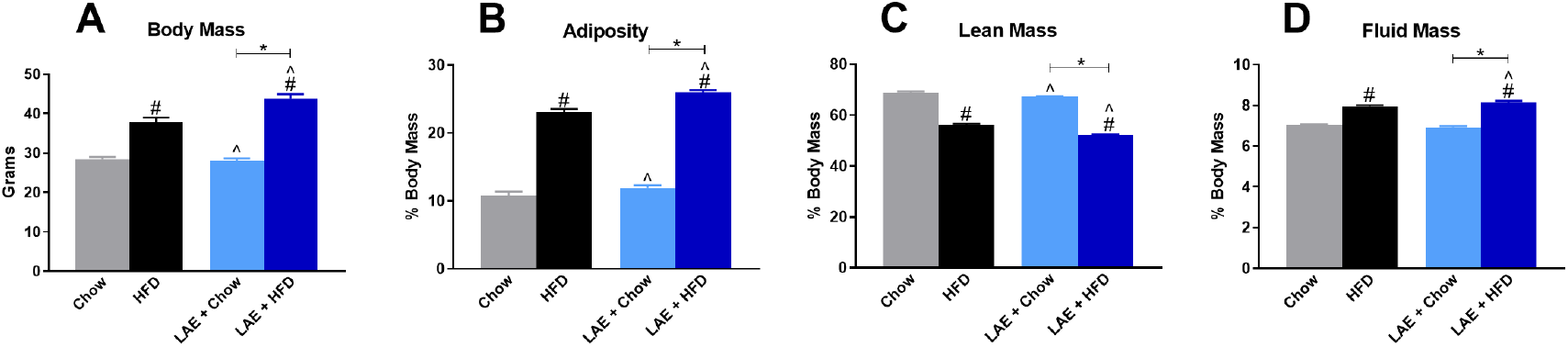
Limited Access EtOH (LAE) consumption worsens HFD-induced changes to body mass, adiposity, lean mass, and fluid mass. Body composition data are compared to chowand HFD-fed EtOH-naïve mice. **A)** Body mass of control mice and LAE mice prior to metabolic testing. Percent body mass of **B)** adiposity, **C)** lean mass, and D) fluid mass. * indicates significant difference between indicated groups, # indicates significant difference from Chow, ^ indicates significant difference from HFD, as determined by one-way ANOVA with Bonferroni’s post-hoc test, p<0.05.

**Fig 9.**
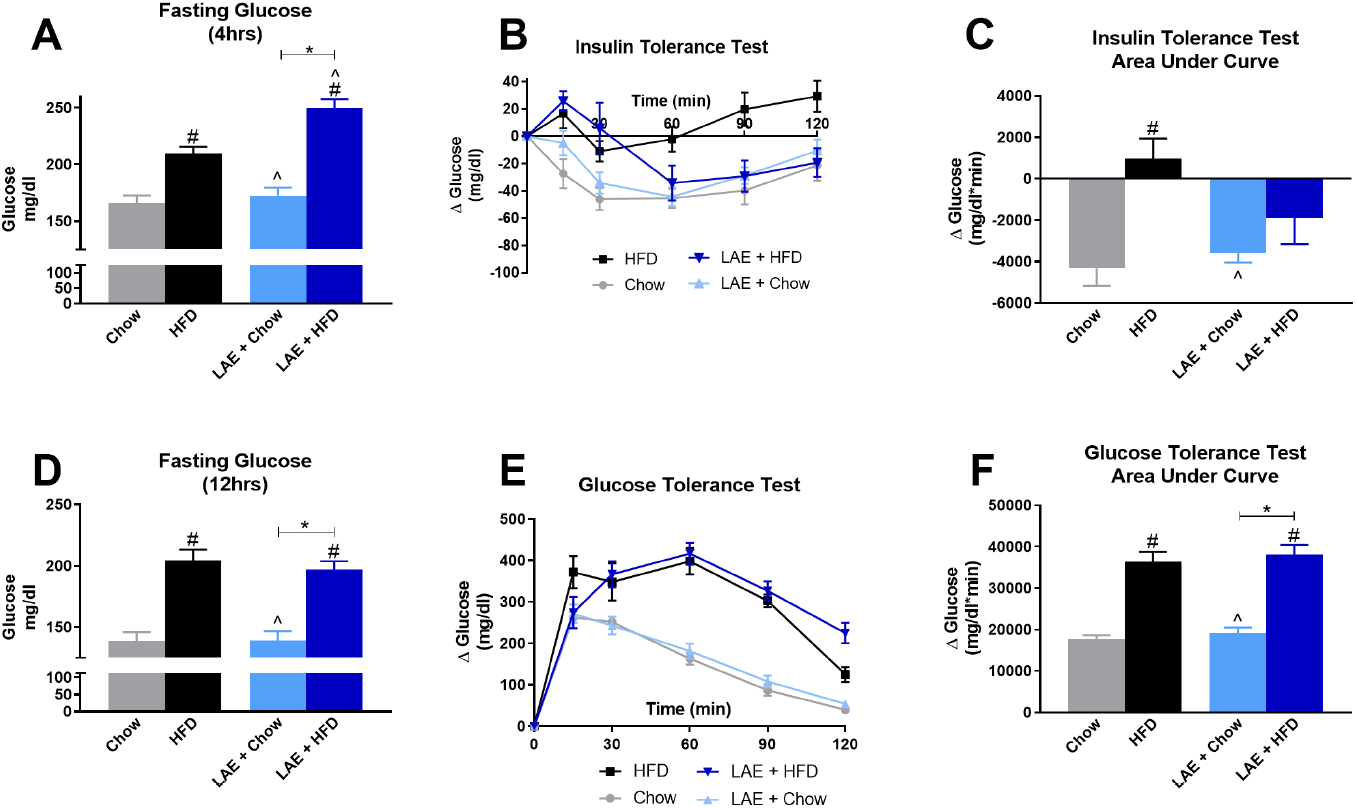
Limited Access EtOH (LAE) does not improve insulin sensitivity or glucose tolerance in HFD-fed mice. **A)** 4-hour fasting glucose levels prior to insulin tolerance test. **B)** Change in blood glucose levels over time following insulin injection; data are normalized to 0 at baseline. **C)** Area under the curve for change in blood glucose levels during insulin tolerance test. **D)** 12-hour fasting glucose levels prior to glucose tolerance test. **E)** Change in blood glucose levels over time following dextrose injection; data are normalized to 0 at baseline. **F)** Area under the curve for change in blood glucose levels during glucose tolerance test. One-way ANOVA; * indicates significant difference between indicated groups, # indicates significant difference from Chow, ^ indicates significant difference from HFD, as determined by one-way ANOVA with Bonferroni’s post-hoc test, p<0.05

**Fig 10.**
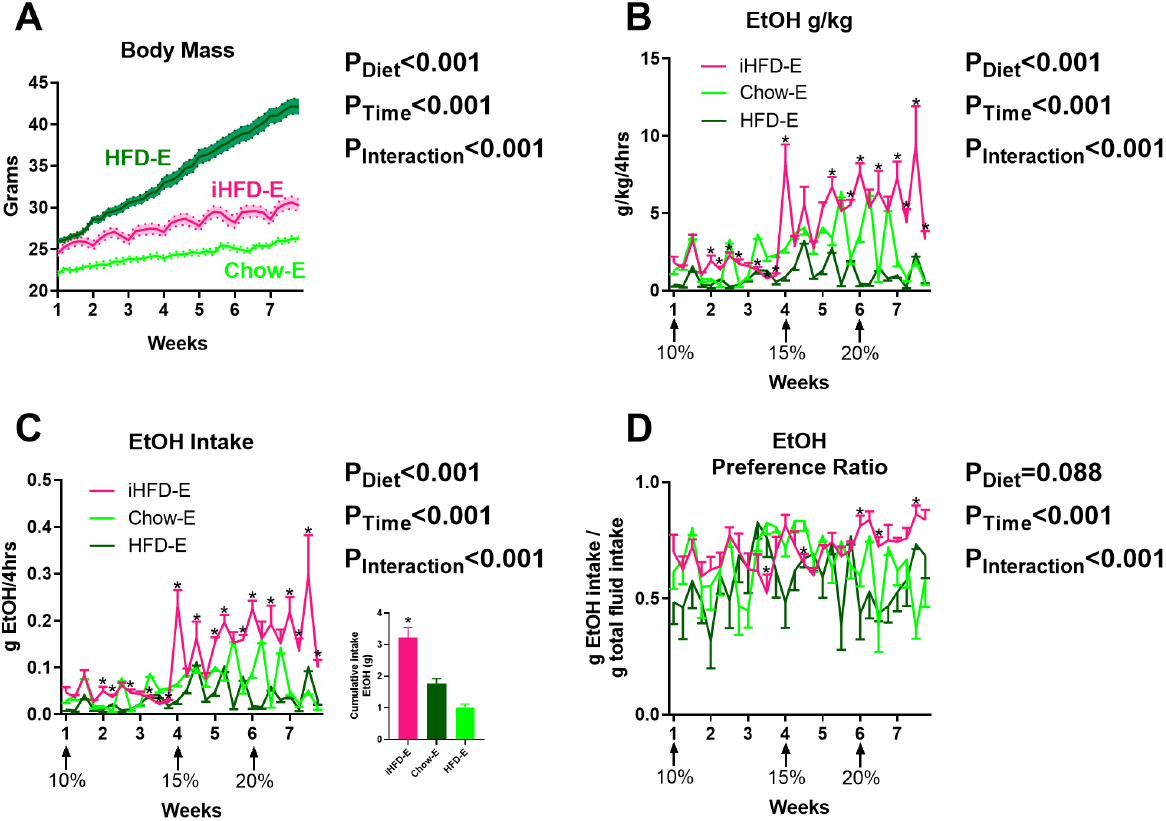
Intermittent HFD effects on limited access to EtOH, “iHFD-E” model. **A)** Time course of body mass changes by group during drinking period (n=10/group). iHFD-E increases g/kg **(B)** and total g/EtOH consumed over the course of the study **(C)** and modestly alters preference **(D)**. *indicates significant differences between groups on individual EtOH access sessions as determined by Bonferroni post-hoc analysis; p<0.05.

**Fig 11.**
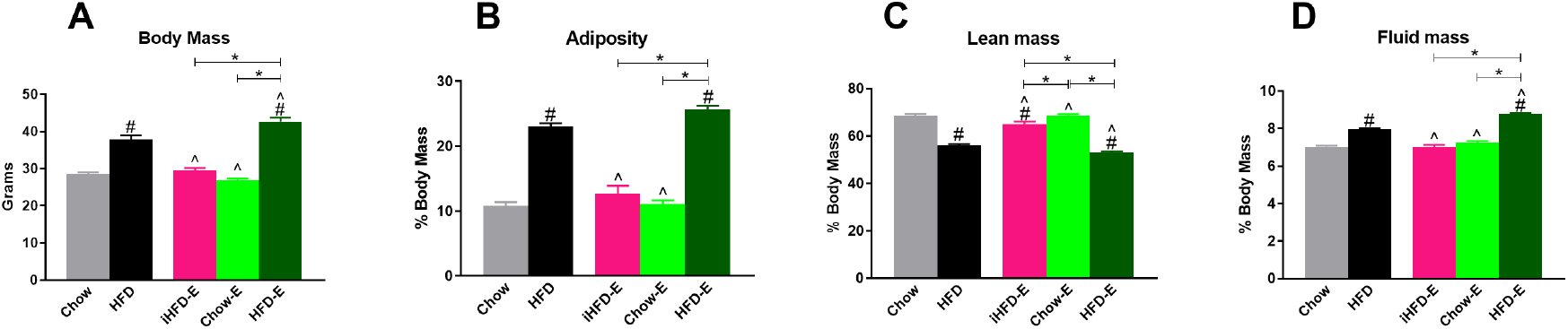
Intermittent HFD does not alter body mass, adiposity, lean mass, or fluid mass. Body composition data are compared to chow- and HFD-fed EtOH-naïve mice. **A)** Body mass of control mice and LAE mice prior to metabolic testing. Percent body mass of **B)** adiposity, **C)** lean mass, and **D)** fluid mass. * indicates significant difference between indicated groups, # indicates significant difference from Chow, ^ indicates significant difference from HFD, as determined by one-way ANOVA with Bonferroni’s post-hoc test, p<0.05.

**Fig 12.**
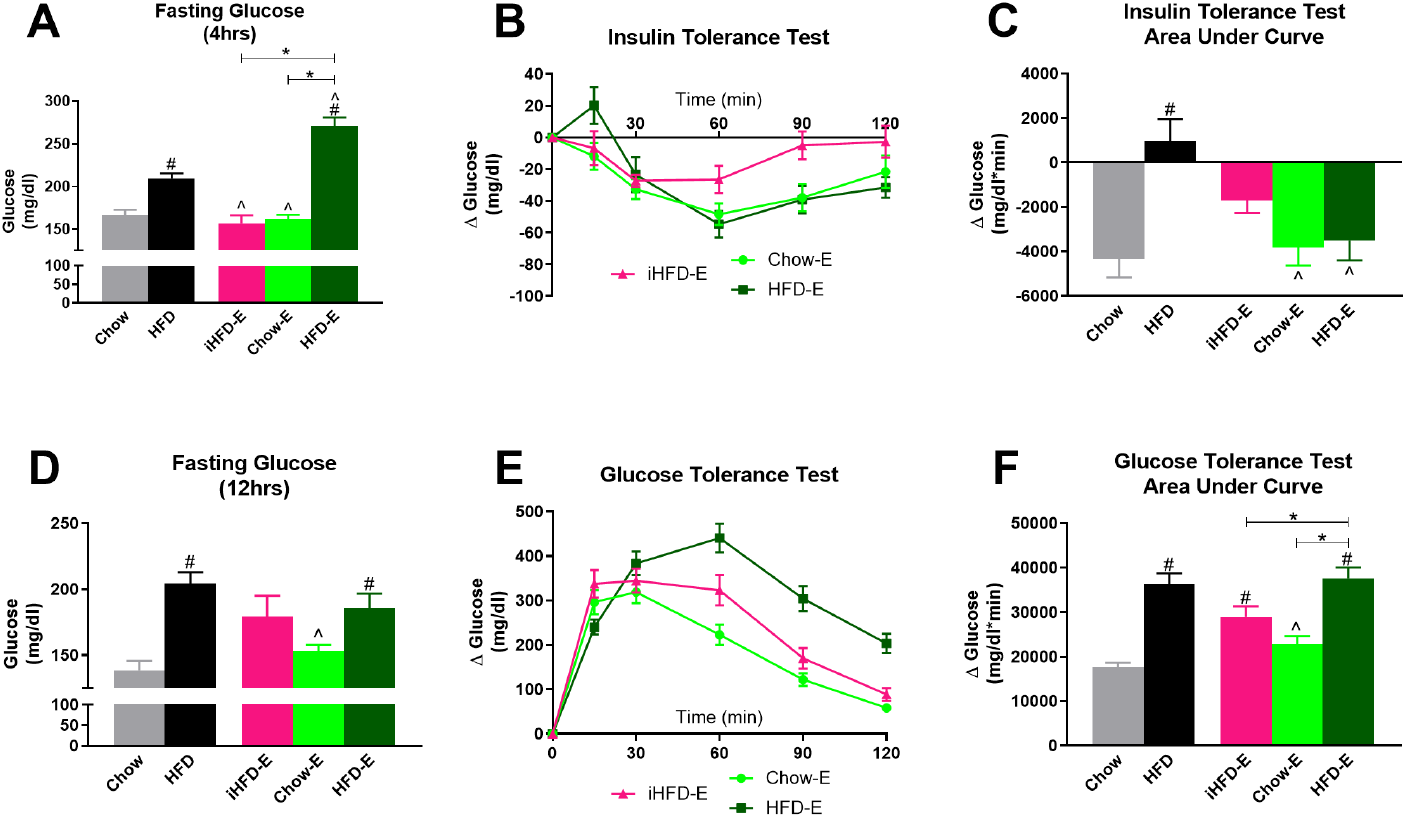
Intermittent HFD promotes insulin resistance and glucose intolerance. **A)** 4-hour fasting glucose levels prior to insulin tolerance test. **B)** Change in blood glucose levels over time following insulin injection; data are normalized to 0 at baseline. **C)** Area under the curve for change in blood glucose levels during insulin tolerance test. **D)** 12-hour fasting glucose levels prior to glucose tolerance test. **E)** Change in blood glucose levels over time following dextrose injection; data are normalized to 0 at baseline. **F)** Area under the curve for change in blood glucose levels during glucose tolerance test. * indicates significant difference between indicated groups, # indicates significant difference from Chow, ^ indicates significant difference from HFD, as determined by one-way ANOVA with Bonferroni’s post-hoc test, p<0.05.

## Discussion

The current study aimed to examine the impact of HFD on EtOH intake in distinct scheduled access periods. Overall, the findings indicate that *ad libitum* HFD access only altered EtOH intake when EtOH intake was *ad libitum* (UAE+HFD mice, Experiment 1). *Ad libitum* HFD however, did not significantly alter EtOH intake when EtOH access was limited (LAE+HFD, Experiment 2; HFD-E, Experiment 3). Intermittent HFD scheduling (iHFD-E, Experiment 3) induced binge eating behaviors that resulted in escalated EtOH intake on days in which HFD was not available. These findings suggest that scheduling access is an important factor in determining the role of HFD on EtOH intake, which has often been overlooked in previous studies on HFD and EtOH interactions. Furthermore, these findings suggest that HFD intake binge behaviors in a limited access model can transfer towards binge EtOH intake behaviors.

Following HFD and EtOH intake periods, body composition and glucose and insulin function was then assessed in these mice. Although UAE+HFD mice in Experiment 1 had lower EtOH intake, this moderate level of EtOH intake did not alter body mass, adiposity, lean mass, fluid mass, or insulin sensitivity compared to mice given HFD for the same amount of time without EtOH. LAE+HFD mice in Experiment 2 had similar levels of EtOH intake compared to LAE+Chow mice, and this level of EtOH intake appears to synergize with HFD consumption to promote increased body mass, adiposity, lean mass, or fluid mass, with similar levels of insulin resistance and glucose intolerance compared to HFD control mice. Similar results were found in the HFD-E mice in Experiment 3, where EtOH consumption further enhanced HFD-induced changes in body composition and fasting glucose levels. Interestingly, while iHFD-E mice in Experiment 3 remained lean and had body composition similar to Chow control mice, iHFD-E mice had insulin and glucose sensitivity similar to HFD mice. This suggests even moderate amounts of HFD consumption in the face of enhanced EtOH intake may synergize to produce metabolic deficits even with a lack of body composition changes. Overall, the total of the findings here indicate that EtOH access schedules are critical in mediating the impact of HFD on EtOH intake, while patterned EtOH intake in the face of HFD may synergize to produce more profound metabolic disturbances than HFD alone.

Our finding that *ad libitum* HFD access decreases *ad libitum* EtOH intake (UAE+HFD group) is a common finding in the literature in both male and female rodents (Feng *et al*, 2012; Gelineau *et al*, 2017; Sirohi *et al*, 2017a, 2017b). This may be due to a number of factors. One possibility is that rodents find HFD a rewarding food choice and prefer this to the potential rewarding effects of EtOH. Although not directly assessed here, animals exposed to HFD typically undergo an initial hyperphagic response (Hariri and Thibault, 2010), suggesting this diet has some rewarding value leading to escalation of intake at least short-term. Further research supports this hypothesis by showing that HFD exposure can alter dopamine activity in the nucleus accumbens (Fordahl *et al*, 2016; Rada *et al*, 2012), ventral tegmental area (Valdivia *et al*, 2015), and alter neuronal signaling in other key brain regions mediating reward processing (Barson *et al*, 2012; Sharma *et al*, 2013; Valdivia *et al*, 2014). The current finding that UAE+HFD mice have a lower EtOH preference than UAE+Chow mice further supports this hypothesis. We only examined 10% EtOH intake in the UAE cohort, however, and it will be important to examine the preference and intake of EtOH at higher concentrations such as was performed in the LAE cohort. Another possibility is that EtOH metabolism and/or clearance may have been altered in UAE+HFD mice due to changes in body composition (increased adiposity and decreased lean mass) when compared with UAE+Chow mice (Feldstein, 1978; Reed and Kalant, 1977). C57Bl/6J mice typically consume enough EtOH to reach pharmacologically relevant BEC levels in relatively short access periods (~ two hours) (Becker and Lopez, 2004). C57Bl/6J mice have also been shown to have numerous drinking bouts when EtOH is provided over extended time periods (Risinger *et al*, 1998). Therefore, although not directly examined here, it is plausible that the UAE+Chow mice may have had numerous bouts of drinking over the 24-hour access period. If UAE+HFD mice had similar initial bouts but prolonged EtOH clearance/metabolism due to changes in adiposity or lean mass, then they may not seek as much EtOH in subsequent bouts to maintain pharmacologically relevant BECs, thus lowering their total EtOH consumption over the 24-hour access period. This possibility will be fully addressed in future studies.

Extending the hypothesis that HFD has a higher relative reward value than EtOH, LAE+HFD mice had significantly lower EtOH preference compared to controls. This effect appeared to be more pronounced at 10% EtOH, with less differences in EtOH preference at 15% and 20% EtOH. This finding may suggest that the rewarding value of 10% EtOH was lower in HFD exposed mice across the two models, but that this may be overcome at higher EtOH doses. Overall though, 10% EtOH consumption was low in the LAE model but increased significantly at higher concentrations, suggesting that the overall reward value of 10% EtOH may have generally been low when given in a 4-hour access period. Therefore, caution must be used when directly comparing the UAE and LAE models given the differences in EtOH access schedules (24 vs 4 hrs, continuously vs intermittently). Importantly, findings from the LAE model suggest that HFD does not greatly alter EtOH intake when EtOH access is limited, regardless of EtOH reward value. This is an important consideration given the previous preclinical literature indicating a reduction in EtOH intake by HFD, but numerous clinical findings indicate that EtOH stimulates HFD intake and vice versa (Breslow *et al*, 2013; Caton *et al*, 2004; Feng *et al*, 2012; Gelineau *et al*, 2017; Piazza-Gardner and Barry, 2014; Sirohi *et al*, 2017a, 2017b). Intriguingly, mice that had HFD on a limited, intermittent schedule (iHFD-E mice) developed binge eating patterns that appeared to trigger increased EtOH consumption in a limited access schedule. This model may better recapitulate the clinical findings above that HFD and EtOH may stimulate over-consumption of both reward modalities. Given the shared neurocircuitry involved in reward value of palatable diets and EtOH (Kenny, 2011), it is possible that binge HFD consumption may have sensitized this shared neurocircuitry to produce higher EtOH intake levels. Examination of this hypothesis will be of great interest in future studies.

Extensive previous research has shown moderate EtOH consumption improves insulin sensitivity in both clinical (Traversy and Chaput, 2015) and preclinical settings (Hong *et al*, 2009). Therefore, it was surprising that UAE+HFD mice, which consumed a moderate amount of EtOH on a per day basis for this mouse strain, had insulin resistance of a similar magnitude to HFD mice without EtOH intake history. The level of HFD-induced insulin resistance in control HFD mice here is similar to our previous research (Loloi *et al*, 2018; Williams *et al*, 2016). Consistent with the finding for insulin resistance, 4-hour fasting glucose in UAE+HFD mice was elevated to a similar degree as observed in the HFD mice. The reason for the lack of replication between our current study and previous research indicating EtOH consumption improves insulin sensitivity in HFD exposed animals is unclear, but previous research does indicate that twice daily intra-gastric EtOH exposure was more beneficial to improve insulin sensitivity in HFD-fed rats than continuous free access intake, even when total daily EtOH dosage (5g/kg) was accounted (Feng *et al*, 2012). This was due to differences in peak plasma EtOH concentrations of the different modes of exposure. Since daily EtOH intake in the UAE+HFD mice was generally between 2-3g/kg per day, this level of EtOH intake may not have been high enough to produce beneficial effects on insulin sensitivity.

In contrast to this hypothesis, LAE+HFD mice, which had higher EtOH intake levels than UAE+HFD mice (~5g/kg per day over the last three weeks of the study) did not have improved insulin sensitivity compared to HFD controls. Body composition and glucose tolerance was also drastically disturbed in the LAE+HFD. Specifically, LAE+HFD mice had increased body mass, adiposity, and fluid mass, with decreased lean mass compared to EtOH-naïve HFD mice. In addition, these mice had elevated 4-hour fasting glucose levels and were glucose intolerant. Such changes in body composition and insulin and glucose function were not seen in LAE+Chow mice, suggesting that the enhanced metabolic disturbances in LAE+HFD were not just additive effects of EtOH on top of those seen in HFD mice, but potentially a synergistic effect. HFD-E mice in the Experiment 3 also had similar, and potentially synergistic, effects of EtOH and HFD consumption on body composition and insulin and glucose function. iHFD-E mice, which drank significantly more EtOH than HFD-E mice but did not gain as much body mass, also had trends toward insulin insensitivity and had significant glucose intolerance. It is possible that such disturbances would have continued to worsen over longer exposure periods in this model, further suggesting synergistic HFD and EtOH interactions. Although the current data cannot speak to the precise mechanism of this potential synergistic action, previous findings indicate that EtOH can greatly impact glucose metabolism by a variety of mechanisms including impairments in intestinal glucose absorption, endogenous pancreatic insulin secretion, glucose effectiveness, and counter-regulatory responses (Steiner *et al*, 2015). Future studies using sophisticated hyperinsulinemic-euglycemic clamp with isotropic tracer methods will be needed to identify tissue-specific disturbances in insulin and glucose action as seen in the LAE+HFD and iHFD-E models.

Overall, the models of combined EtOH and HFD consumption described here point to little benefit of EtOH in the face of metabolic dysfunction. The concept that moderate EtOH drinking has beneficial health effects has come under increased scrutiny in the past few years (Griswold *et al*, 2018) and brings back into debate the potential interactive role of EtOH and HFD in the development of metabolic diseases, such as Type II diabetes. Indeed, given the well-established roles of EtOH and HFD as individual risk factors for the development of metabolic disturbances, and the increasing understanding that chronic EtOH and HFD have similar effects on peripheral and central signaling mechanisms, it is surprising that clinical evidence points to moderate EtOH consumption as a mitigating factor in HFD-induced metabolic disturbances. The findings here suggest that there are many factors that may influence how EtOH and HFD interact to promote or mitigate metabolic disturbances, such as frequency and duration of EtOH access. It should be noted that several studies report a U- or J-shaped relationship between EtOH and insulin function (Kiechl *et al*, 1996; Lazarus *et al*, 1997; Villegas *et al*, 2004), or that beneficial effects of EtOH may only be seen in those individuals without obesity or insulin resistance (Yokoyama, 2011). Such findings further suggest the need to better examine the interactions of EtOH and HFD on insulin action and glucose tolerance both clinically and pre-clinically while controlling for time course, duration, and frequency of both EtOH and HFD exposures.

